# Dose-specific effectiveness of 7- and 13-valent pneumococcal conjugate vaccines against vaccine-serotype *Streptococcus pneumoniae* colonization: a matched, case-control study

**DOI:** 10.1101/532085

**Authors:** Joseph A. Lewnard, Noga Givon-Lavi, Ron Dagan

**Affiliations:** Division of Epidemiology, School of Public Health, University of California, Berkeley, Berkeley, California, United States; Ben-Gurion University of the Negev, Be’er Sheva, Israel

**Keywords:** *Streptococcus pneumoniae*, pneumococcal conjugate vaccine (PCV), vaccine effectiveness, carriage, vaccine schedule, 2+1

## Abstract

**Background:** Reduced-dose pneumococcal conjugate vaccine (PCV) schedules are under consideration in countries where children are currently recommended to receive three PCV doses. However, dose-specific PCV effectiveness against vaccine-serotype colonization is uncertain.

**Methods:** From 2009-2016, we conducted surveillance of pneumococcal carriage in southern Israel, where PCV is administered at ages 2, 4, and 12 months (2+1 schedule). We obtained nasopharyngeal swabs and vaccination histories from 4245 children ages 0-59 months without symptoms of diseases that could be caused by pneumococci. In a case-control analysis, we measured protection against vaccine-serotype colonization as one minus the matched odds ratio for PCV doses received.

**Results:** At ages 5-12 months, a second PCV7/13 dose increased protection against PCV7-serotype carriage from –23.6% (95%CI: –209.7-39.1%) to 27.1% (–69.2-64.5%), and a second PCV13 dose increased protection against carriage of all PCV13 serotypes from –54.8% (–404.3-39.1%) to 23.4% (– 128.5-67.1%). At ages 13-24 months, a third PCV7/13 dose increased protection against PCV7-serotype carriage from 32.4% (–8.4-58.0%) to 74.1% (58.4-84.6%), and a third PCV13 dose increased protection against carriage of all PCV13 serotypes from –50.0% (–194.0-42.7%) to 49.7% (15.8-83.3%). On average, each PCV13 dose conferred 37.7% (7.0-61.8%) greater protection against carriage of serotypes 1, 5, 6A, 7F, and 19A than carriage of serotype 3. PCV13-derived protection against carriage of serotypes 1, 5, 6A, 7F, and 19A was equivalent to PCV7/13-derived protection against carriage of PCV7 serotypes.

**Conclusions:** In a setting implementing a 2+1 PCV schedule, protection against vaccine-serotype colonization is sustained primarily by the third dose.

## INTRODUCTION

Commensal carriage of *Streptococcus pneumoniae* in the nasopharynx is the source of transmission and a precursor to pneumococcal meningitis, bacteremia, pneumonia, and otitis media.^1^ Pneumococcal conjugate vaccines (PCVs) targeting 7, 10, and 13 of the 96+ pneumococcal serotypes have been licensed and introduced into pediatric immunization programs of 59 countries throughout the world.^2^ Implementation of pediatric PCV programs has reduced incidence of invasive pneumococcal disease (IPD) caused by vaccine-serotype pneumococci among vaccinated children, and unvaccinated children and adults experiencing indirect protection.^3^ However, the high cost of PCVs strains the budgets of national healthcare programs and charities.^2^ Recently, this expense has drawn the continuation of PCV10/13 immunization programs in certain middle-income countries (LMICs) into question as they phase out full support for PCVs from GAVI, the vaccine alliance.^4^

Most countries implement a two- or three-dose primary PCV series in the first year of life, with or without a booster dose at age 12 months (3+1, 2+1, and 3+0 primary+booster schedules). The possibility of sustaining protection with fewer doses has thus attracted interest as a cost-saving measure. A recent trial in the UK demonstrated non-inferior immunogenicity of a 1+1 PCV13 series, relative to the country’s 2+1 schedule.^5^ However, the ability of reduced-dose schedules to sustain the population-wide impact achieved by PCV programs to date depends on effectiveness against vaccine-serotype colonization.^6^ To date, the effectiveness of each PCV dose on vaccine-serotype colonization has not been systematically assessed.^3,7,8^

Because many children do not receive PCVs according to recommended schedules, observational studies comparing outcomes among children with differing vaccination histories can provide evidence of the potential effectiveness of alternative dosing schemes.^9^ Such studies also provide crucial insight into real-world vaccine performance. Long-term population-based surveillance in southern Israel has provided a window into pneumococcal colonization among children in this setting amid PCV7/13 implementation.^10^ To guide scheduling considerations, we estimated PCV direct effects against vaccine-serotype colonization according to doses-received-for-age in an individually matched, case-control study.

## METHODS

### Setting

Prospective surveillance of pneumococcal carriage is undertaken at Soroka University Medical Center (SUMC) in Be’er Sheva, Israel. As the only medical center in Southern Israel (the Negev District), SUMC is where >90% of all births and >90% of all pediatric emergency room visits and hospitalizations of children <5 years old occurs, enabling population-based studies. The surrounding region is inhabited by socioeconomically-distinct Jewish and Bedouin (Muslim) populations. Despite their geographic proximity, Jewish and Bedouin children have limited contact. The Bedouin population is transitioning from a semi-nomadic lifestyle to permanent settlements; in comparison to Jewish communities, Bedouin communities are distinguished by lower household income, higher birthrates and larger family sizes, and poorer health status, despite receiving care at the same facilities.^11^ Prior to PCV7/13 implementation, the region’s Bedouin and Jewish populations resembled those of lower- and higher-income countries, respectively, in pneumococcal carriage and disease.^12^

Uptake of PCVs in southern Israel has been described previously.^10^ A 2+1 PCV7 schedule (with doses administered at ages 2, 4, and 12 months) was introduced in July, 2009, with a catch-up campaign among children under 2 years of age. Beginning in November, 2010, PCV13 replaced PCV7 without additional catch-up. We illustrate PCV7/13 coverage trends by age in **Figure 1**.

**Figure 1:**
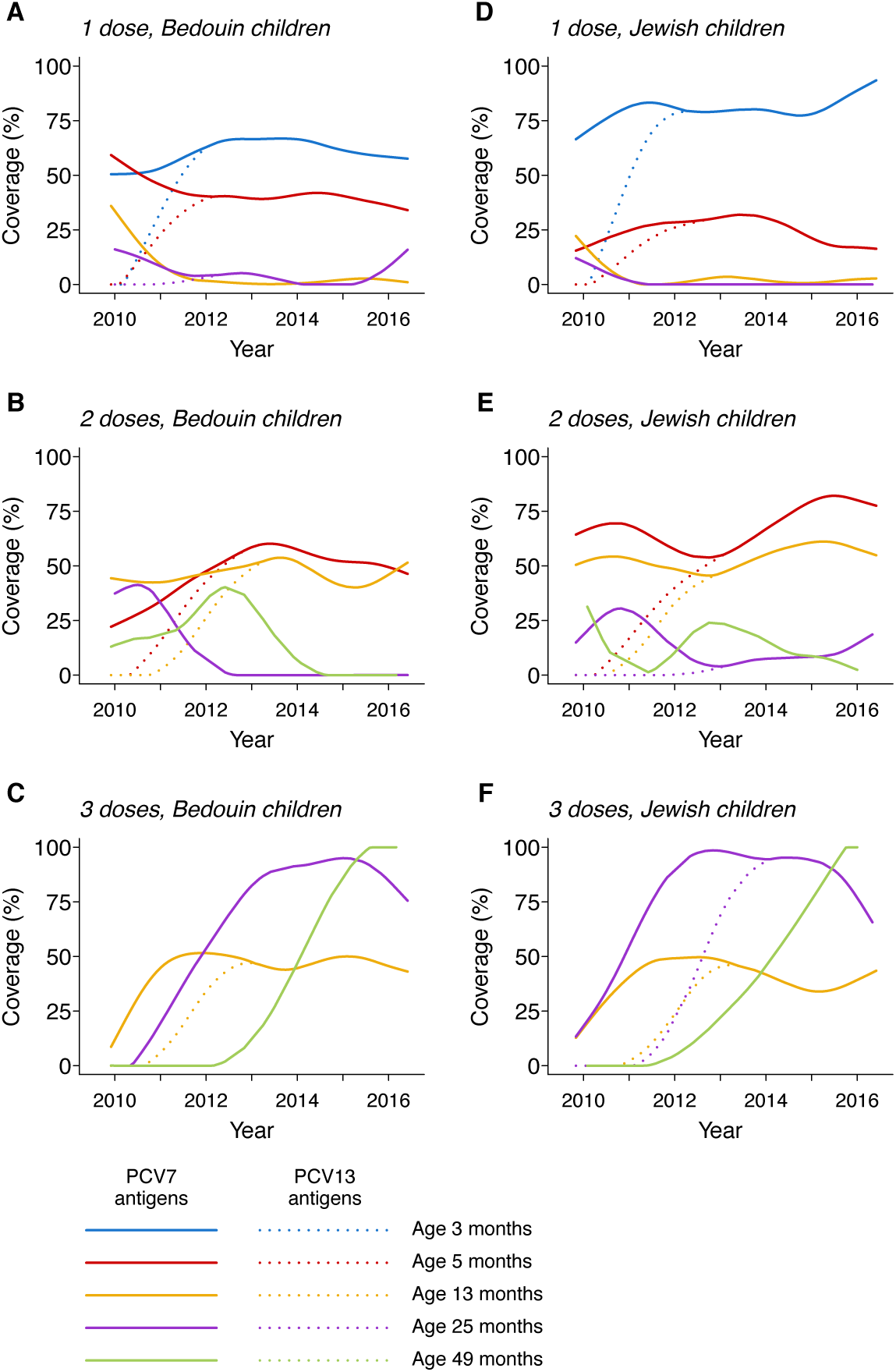
Vaccine coverage in the study population. We illustrate trends in axsge-specific coverage of immunization against PCV7-type (solid lines) and PVC13-type (dotted lines) antigens in the Bedouin and Jewish populations, among all children enrolled in the study. Israel added PCV7 to the pediatric vaccination schedule in late 2009, using a 2+1 dosing scheme, with a catch-up campaign for children under 2 years old. Substitution of PCV13 for PCV7 began in November, 2010. We fitted trends to data using smoothing splines with 5 knots.

### Surveillance study

Surveillance of pneumococcal carriage among Jewish and Bedouin children <5 years old has been undertaken at SUMC since PCV7 introduction in 2009; each working day, a nasopharyngeal swab is obtained from the first four Jewish and first four Bedouin children presenting for any reason to the pediatric emergency department, who are residents of the Negev region, and whose parents provide informed consent. Previous pneumococcal vaccination (number of PCV7 and PCV13 doses) were ascertained from children’s medical records.^10^

Nasopharyngeal samples were obtained via dacron-tipped swabs and plated within 16 hours on 5% sheep blood/5.0μg gentamicin agar media, and incubated for 48 hours at 35°C. Pneumococcal identification was based on alpha-hemolysis and optochin inhibition, and confirmed by slide agglutination. One colony per plate was selected for serotyping by the Quellung reaction (Statens Seruminstitut, Copenhagen, Denmark), as described previously.^10^

### Case-control framework

We measured vaccine effectiveness in a case-control framework comparing the prevalence of prior PCV receipt among “case” children, who experienced the target endpoint of vaccine-serotype colonization, versus “control” children, who were sampled from the eligible study population at random. While previous studies of pneumococcal vaccine effectiveness have defined individuals with nonvaccine-serotype infection as controls,^13,14^ PCV increases individuals’ risk of nonvaccine-serotype carriage,^15–17^ making such methods inappropriate for studies of colonization. We detail the statistical rationale and procedure for use of randomly-sampled children as controls in the **Supporting information**.

To counteract potential confounding, we matched case and control children according to the following attributes: ethnicity (Jewish or Bedouin), age (within 1 month of the case child’s age), visit timing (within 2 months of the case child’s visit), and recent antibiotic receipt (receiving any antibiotic within 1 month before the visit). To ensure analyses addressed PCV effects on carriage and not pneumococcal disease syndromes, we included children in the case-control study whose diagnoses did not include otitis media, pneumonia, influenza, lower or upper respiratory infection, conjunctivitis, bacteremia/sepsis, or meningitis.

### Estimation of vaccine effectiveness

#### Endpoints

We assessed the effectiveness of primary series doses by comparing prevalence of receipt of 0, 1, or 2 PCV doses among case and control children ages 5-12 months. We assessed the long-term effectiveness of the third dose by comparing prevalence of receipt of 0, 2, and 3 doses among case and control children ages 13-24 months and 13-59 months.

We assessed protection for several vaccine exposures and carriage endpoints. First, we assessed protection against carriage of serotypes 4, 6B, 9V, 18C, 19F, and 23F (PCV7-targeted serotypes) conferred by differing numbers of PCV7 or PCV13 doses. We next assessed protection against PCV7-targeted serotypes conferred by PCV7 in analyses excluding children who received any PCV13 doses, and protection against PCV7-targeted serotypes conferred by PCV13 in analyses excluding children who received any PCV7 doses. Last, we assessed PCV13 effectiveness against carriage of all PCV13 serotypes, and against carriage of the six serotypes targeted only by PCV13 (1, 3, 5, 6A, 7F, and 19A; +6PCV13 serotypes), in analyses excluding children who received any PCV7 doses.

#### Statistical analysis

We conducted statistical inference in a resampling framework to account for the permutation distribution of case-control matching assignments, as detailed in the **Supporting information**. For each iteration, we matched each case to six controls and estimated dose-specific PCV effectiveness as one minus the matched odds ratio for receipt of differing numbers of PCV doses. We estimated matched odds ratios using conditional logistic regression models, including matching strata. We generated distributions of VE estimates across 2,000 independent iterations of this assignment and estimation scheme.

### Secondary analyses

#### Differential protection by ethnicity

We also sought to assess whether protection differed for serotypes targeted by PCV7, those targeted by PCV13, and for serotype 3, as previous studies have suggested.^14^ Because analyses were underpowered for comparison of age- and dose-specific effectiveness across strata, we used conditional logistic regression to compare the degree of protection indicated by the continuous trend in the matched odds ratio for receipt of one to three doses, as compared to zero doses. Matched sets were assembled according to the approach described above for VE estimation.

We examined differences in protection according to two measures. First, we calculated the difference in the average per-dose reduction in odds of the various endpoints (here defined *i* and *j*) as

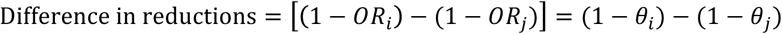

for *θ*_*i*_ regression coefficients indicating the average reduction, per PCV dose received, in odds of carrying serotype(s) *i* (see **Supporting information**).

We also estimated the average marginal increase in protection against each outcome, relative to other outcomes, suggested by the trends in slopes. For the relative degree of protection against serotype(s*) i* versus protection against serotype(s) *j*, this measure was

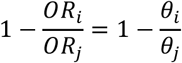

for *θ*_*i*_ as defined above.

#### Differential protection by serotype

We used the same approach to assess differences in protection conferred against each endpoint (carriage of PCV7 serotypes, PCV13 serotypes, +6PCV13 serotypes, serotypes 1, 5, 6A, 7F, and 19A, and serotype 3) in Bedouin and Jewish children. We calculated the difference in protection between the two populations as

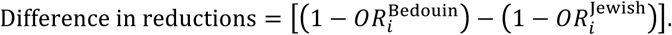

## RESULTS

### Enrollment

Between November, 2009 and June, 2016, nasopharyngeal swabs were processed from 5780 Bedouin and 4386 Jewish children ages 0-59 months, among whom 2436 Bedouin children and 1809 Jewish children were not diagnosed with otitis media, pneumonia, influenza, lower or upper respiratory infection, conjunctivitis, bacteremia/sepsis, or meningitis. All children ages 0-24 months at the beginning of the study were eligible for routine or catch-up PCV7 receipt (**Figure 1**); coverage with PCV13 antigens increased with subsequent birth cohorts according to eligibility for 2+1 dosing.

In total, 1656 eligible Bedouin children (47.6%) and 990 eligible Jewish children (39.2%) carried pneumococci (**Table 1**). While PCV7 and PCV13 serotypes comprised over half of all pneumococcal carriage at the beginning of the study period, non-vaccine serotypes comprised an increasing proportion of all carriage over time (**Figure 2**), with serotypes 15B/C and 16F showing the highest prevalence individually (**Figure 3**). Whereas prevalent PCV-targeted serotypes such as 6A, 6B, and 9V were nearly eliminated after PCV7/13 rollout, serotypes 3, 14, 19A, and 19F saw limited, if any, reductions in carriage prevalence.

**Table 1:**
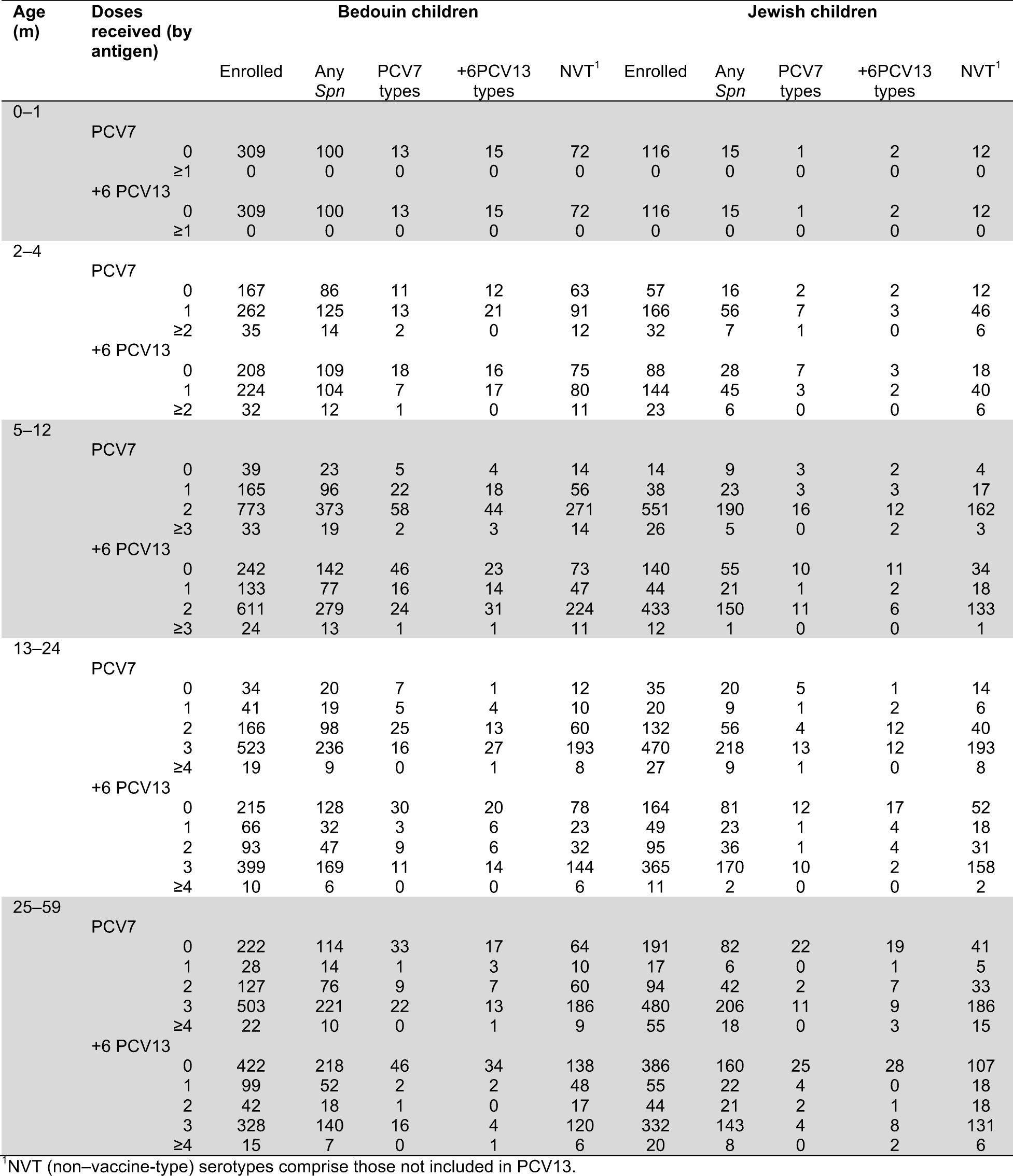
Child-visits by age, carrier status, and vaccination history.

**Figure 2:**
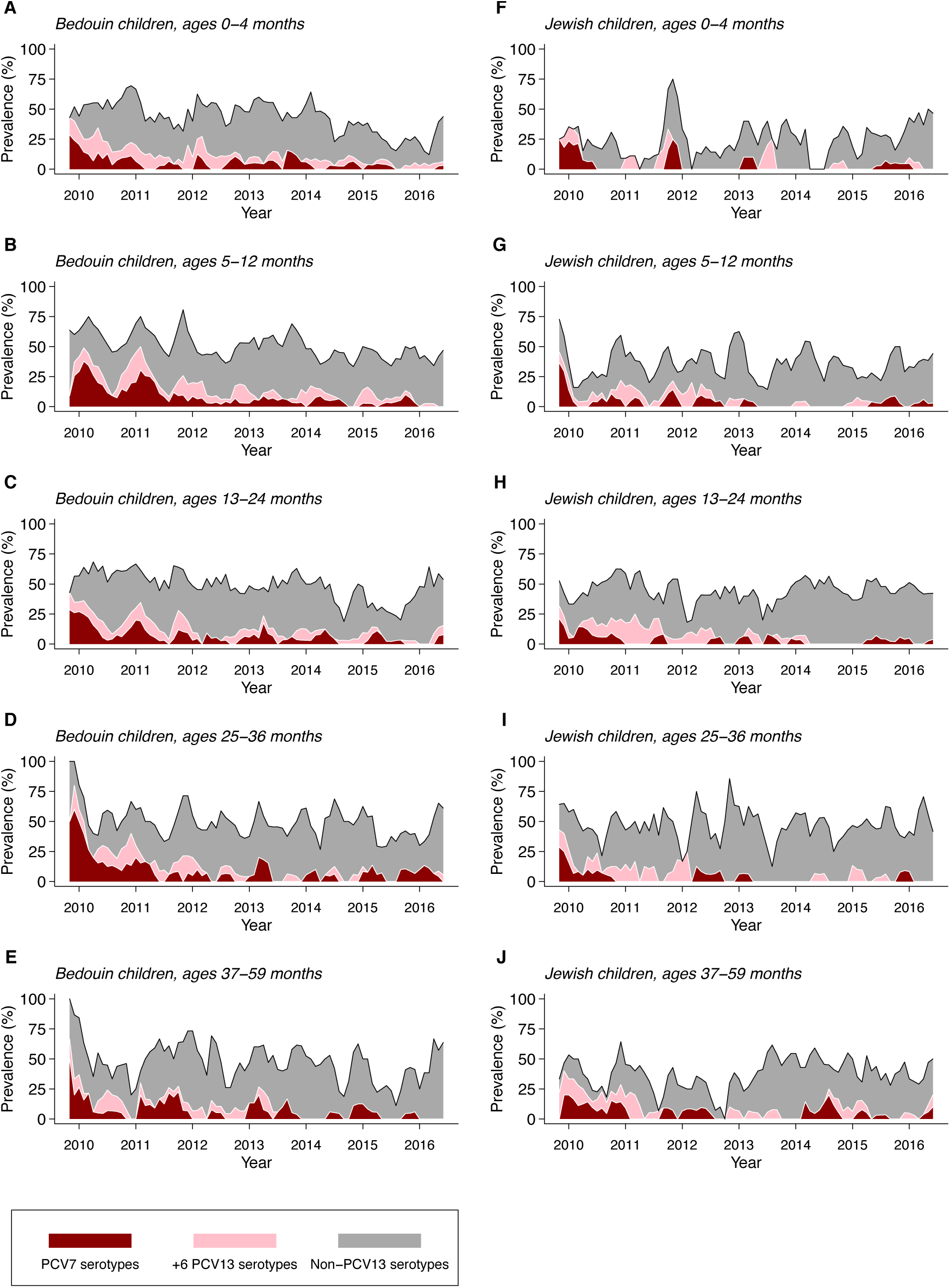
Pneumococcal carriage in the study population. We illustrate pneumococcal carriage prevalence by age, ethnicity, vaccine coverage, and over time (calculated as a 3-month moving average within each age group). Although non-PCV13 serotype carriage largely offset reductions in carriage of vaccine-targeted serotypes, carriage of serotypes targeted by PCV13 persisted as of 2016 in all age groups.

**Figure 3:**
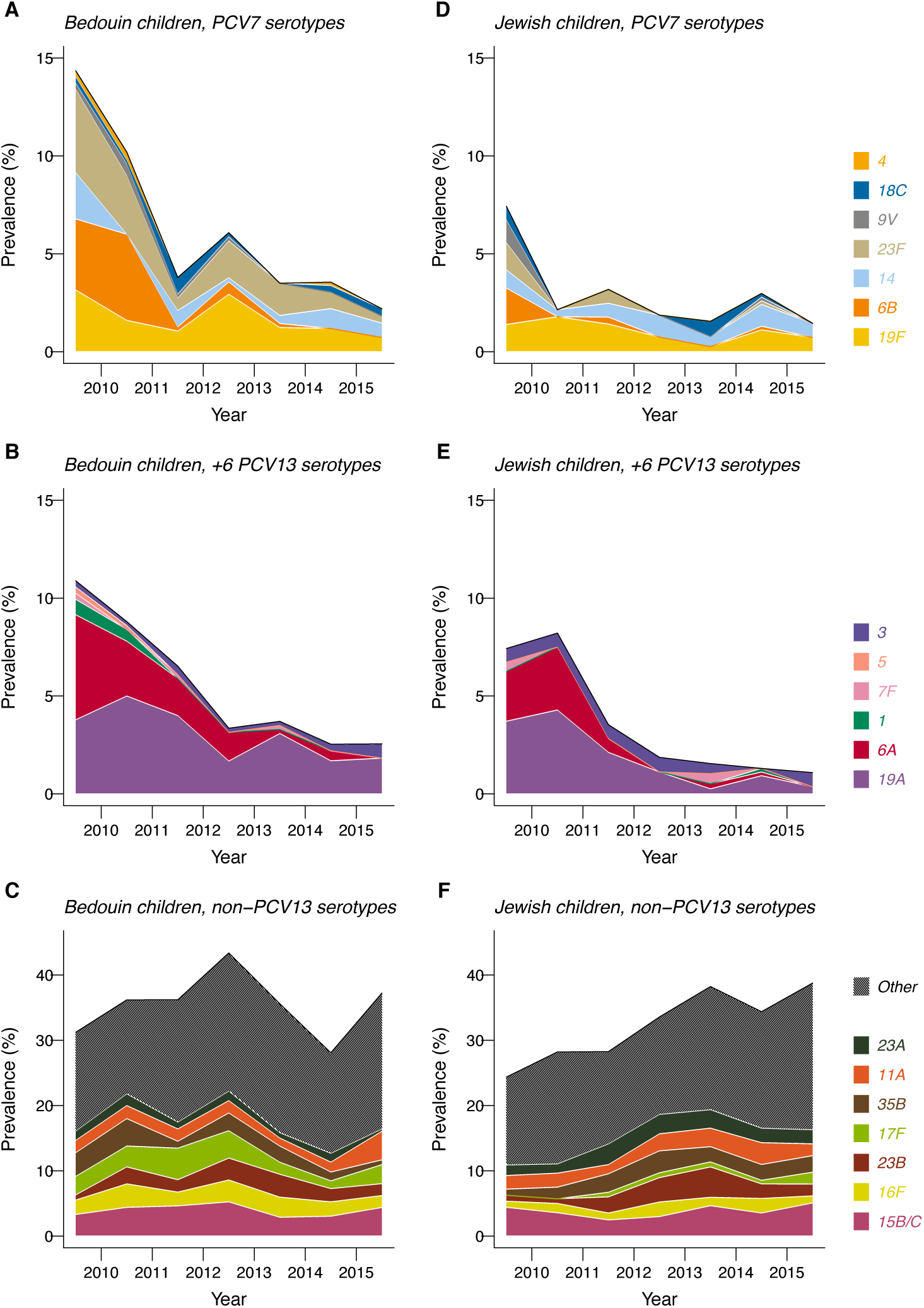
Serotype-specific carriage in the study population. We illustrate prevalence of individual serotypes by ethnicity, year, and vaccine coverage. The predominant PCV13-targeted serotypes persisting after vaccine introduction included 19F, 14, 19A, and 3. The most prevalent non-PCV13 replacement serotypes were 15B/C and 16F.

### Vaccine effectiveness

#### 1- and 2-dose effectiveness against vaccine-serotype carriage at ages 5-12 months

At ages 5-12 months, receipt of one and two PCV7/13 doses conferred –23.6% (–209.7-39.1%) and 27.1% (–69.2-64.5%) protection against PCV7-serotype carriage, respectively, signifying a 41.0% (8.2-60.8%) increase in protection with receipt of the second dose (**Table 2**). Estimates of one- and two-dose effectiveness against PCV7-serotype carriage were 23.1% (–92.9-74.3%) and 30.0% (–65.2-71.0%) for children receiving PCV7 only, representing a 7.1% (–82.4-49.4%) increase in protection following receipt of the second dose. We could not construct sufficient matched sets for estimation of PCV13 effectiveness against carriage of PCV7 serotypes, as prevalence of these serotypes was low in birth cohorts eligible to receive PCV13 in the first year of life.

**Table 2:** Effect of 1 and 2 primary series doses on vaccine-serotype carriage at ages 5–12 months.

| Doses | PCV7/13 |  | PCV13 only |  | PCV7 only |
| --- | --- | --- | --- | --- | --- |
|  | PCV7 serotypes<br>VE (95% CI), % | All PCV13 serotypes<br>VE (95% CI), % | +6 PCV13 serotypes<br>VE (95% CI), % | PCV7 serotypes<br>VE (95% CI), % | PCV7 serotypes<br>VE (95% CI), % |
| 0 | ref. | ref. | ref. | -- | ref. |
| 1 | -23.6 (-209.7, 39.1) | -54.8 (-404.3, 39.1) | 21.1 (-286.2, 79.7) | -- | 23.1 (-92.9, 74.3) |
| 2 | 27.1 (-69.2, 64.5) | 23.4 (-128.5, 67.1) | 50.1 (-121.4, 85.6) | -- | 30.0 (-65.2, 71.0) |
| 1 | ref. | ref. | ref. | -- | ref. |
| 2 | 41.0 (8.2, 60.8) | 49.1 (21.6, 68.8) | 34.4 (-7.0, 62.6) | -- | 7.1 (-82.4, 49.4) |

One and two PCV13 doses conferred –54.8% (–404.3-39.1%) and 23.4% (–128.5-67.1%) protection against all PCV13-serotype carriage, and 21.1% (–286.2-79.7%) and 50.1% (–121.4-85.6%) protection against carriage of +6PCV13 serotypes (**Table 2**). These estimates indicated 49.1% (21.6-68.8%) and 34.4% (–7.0-62.6%) increases in protection against carriage of all PCV13 serotypes and +6PCV13 serotypes, respectively, with receipt of the second PCV13 dose.

#### 2- and 3-dose effectiveness against vaccine-serotype carriage at ages ≥13 months

Receipt of two and three PCV7/13 doses conferred 32.4% (–8.4-58.0%) and 74.1% (58.4-84.6%) protection, respectively, against PCV7-serotype carriage at ages 13-24 months (**Table 3**); the third dose was estimated to provide a 61.6% (46.0-73.4%) increase in protection against PCV7-serotype colonization, over protection sustained with two doses. These estimates of protection in the second year of life were similar to estimates among children receiving PCV7 only, and similar to estimates of protection at ages 13-59 months.

**Table 3:** Effect of 2 and 3 PCV doses on vaccine-serotype carriage after the first year of life.

| Age (m) | Doses | PCV7/13 |  | PCV13 only |  | PCV7 only |
| --- | --- | --- | --- | --- | --- | --- |
|  |  | PCV7 serotypes<br>VE (95% CI), % | All PCV13 serotypes<br>VE (95% CI), % | +6 PCV13 serotypes<br>VE (95% CI), % | PCV7 serotypes<br>VE (95% CI), % | PCV7 serotypes<br>VE (95% CI), % |
| 13–24 |  |  |  |  |  |  |
|  | 0 | ref. | ref. | -- | -- | ref. |
|  | 2 | 32.4 (–8.4, 58.0) | –50.0 (–194.0, 42.7) | -- | -- | 47.2 (8.6, 69.9) |
|  | 3 | 74.1 (58.4, 84.6) | 49.7 (15.8, 83.3) | -- | -- | 76.6 (55.3, 88.9) |
|  | 2 | ref. | ref. | -- | -- | ref. |
|  | 3 | 61.6 (46.0, 73.4) | 65.8 (49.9, 77.6) | -- | -- | 55.8 (25.2, 78.2) |
| 13–59 |  |  |  |  |  |  |
|  | 0 | ref. | ref. | -- | -- | ref. |
|  | 2 | 55.6 (26.5, 75.2) | 45.4 (–47.1, 79.4) | -- | -- | 65.3 (34.8, 86.0) |
|  | 3 | 71.3 (47.1, 84.8) | 55.2 (–10.1, 77.4) | -- | -- | 68.6 (34.1, 88.0) |
|  | 2 | ref. | ref. | -- | -- | ref. |
|  | 3 | 33.5 (–2.4, 60.1) | 12.4 (–64.5, 57.4) | -- | -- | 2.9 (–89.8, 58.8) |

Children who received two and three PCV13 doses experienced –50.0% (–194.0-42.7%) and 49.7% (15.8-83.3%) protection against carriage of all PCV13 serotypes at ages 13-24 months (**Table 3**), with the third dose providing a 65.8% (49.9-77.6%) increase in protection. We could not construct sufficient matched sets to distinguish PCV13-derived protection against PCV7 serotypes or +6PCV13 serotypes.

### Secondary analyses

#### Evidence of serotype-specific protection

On average, each PCV7/13 dose conferred a 27.5% (18.3-36.2%) reduction in odds of carrying PCV7-targeted serotypes (**Table 4**), while each PCV13 dose conferred a 21.2% (5.6-35.8%) reduction in odds of carrying serotypes 1, 5, 6A, 7F, and 19A. In contrast, we estimated the average per-dose reduction in odds of carrying serotype 3 to be –38.9% (–116.7-18.4%).

**Table 4:** Assessing differential protection by serotype.

| Serotype | Vaccine | Doses | Reduction, per dose received (95% CI), % | Difference in reduction (DR) and marginal reduction (MR) per dose received, % |  |  |  |
| --- | --- | --- | --- | --- | --- | --- | --- |
|  |  |  |  | *ref. +5PCV13 serotypes*<br>DR (95% CI)<sup>1</sup> | *MR (95% CI)<sup>2</sup>* | *ref. serotype 3*<br>DR (95% CI)<sup>1</sup> | *MR (95% CI)<sup>2</sup>* |
| PCV7 serotypes | PCV7/13 | 0 | ref. | -- | -- | -- | -- |
|  |  | 1–3 (trend) | 27.5 (18.3, 36.2) | 9.2 (–6.1, 25.5) | 10.5 (–8.6, 27.2) | 65.0 (16.0, 134.8) | 44.7 (17.4, 64.9) |
| Serotypes 1, 5, 6A, 7F, 19A | PCV13 only | 0 | ref. | -- | -- | -- | -- |
|  |  | 1–3 (trend) | 21.2 (5.6, 35.8) | ref. | ref. | 55.7 (6.3, 127.3) | 37.7 (7.0, 61.8) |
| Serotype 3 | PCV13 only | 0 | ref. | -- | -- | -- | -- |
|  |  | 1–3 (trend) | –38.9 (–116.7, 18.4) | –55.7 (–127.3, –6.3) | –69.1 (–161.9, –7.5) | ref. | ref. |

1. DR: Difference in reduction, per dose received, estimated as $(1 - \theta_i) - (1 - \theta_j)$ for a comparison of the magnitude of protection against endpoints $i$ and $j$ .
2. MR: Marginal reduction, per dose received, estimated as $1 - \theta_i / \theta_j$ to quantify the additional protection conferred against endpoint $i$ beyond protection conferred against endpoint $j$ .

Based on these estimates, in comparison to PCV13-derived protection against serotype 3, each PCV13 dose conferred, on average, 37.7% (7.0-61.8%) greater protection against carriage of serotypes 1, 5, 6A, 7F, and 19A, and each PCV7/13 dose conferred 44.7% (17.4-64.9%) greater protection against carriage of PCV7-targeted serotypes (**Table 4**). No difference was evident between trends signifying PCV7/13-derived protection against PCV7-targeted serotypes and PCV13-derived protection against serotypes 1, 5, 6A, 7F, and 19A.

#### Protection among Jewish and Bedouin children

On average, each PCV7/13 dose conferred 29.0% (15.6-41.4%) and 26.1% (15.4-35.1%) reductions in odds of carrying PCV7-serotype pneumococci among Jewish and Bedouin children, respectively (**Table 5**). Similarly, each PCV13 dose conferred 27.0% (11.8-40.9%) and 19.0% (2.4-32.2%) reductions in odds of carrying all PCV13-serotype pneumococci in these populations.

**Table 5:** Assessing differential protection by ethnicity.

| Serotype | Vaccine | Doses | Reduction, per dose received (95% CI), % |  | Difference in reduction per dose received (95% CI), %<br>Bedouin ref. Jewish children |
| --- | --- | --- | --- | --- | --- |
|  |  |  | Jewish children | Bedouin children |  |
| PCV7 serotypes | PCV7/13 | 0 | ref. | ref. | -- |
|  |  | 1–3 (trend) | 29.0 (15.6, 41.4) | 26.1 (15.4, 35.1) | –2.8 (–19.0, 12.8) |
| All PCV13 serotypes | PCV13 only | 0 | ref. | ref. | -- |
|  |  | 1–3 (trend) | 27.0 (11.8, 40.9) | 19.0 (2.4, 32.2) | –8.0 (–28.6, 11.8) |
| +6 PCV13 serotypes | PCV13 only | 0 | ref. | ref. |  |
|  |  | 1–3 (trend) | 37.9 (18.2, 58.2) | 0.0 (–26.3, 20.1) | –37.6 (–69.8, –9.5) |
| Serotypes 1, 5, 6A, 7F, 19A | PCV13 only | 0 | ref. | ref. |  |
|  |  | 1–3 (trend) | 34.8 (11.9, 57.1)<sup>1</sup> | 8.1 (–11.5, 24.5)<sup>2</sup> | –26.1 (–55.5, 0.8) |
| Serotype 3 | PCV13 only | 0 | ref. | ref. |  |
|  |  | 1–3 (trend) | 35.8 (–7.0, 81.7)<sup>1</sup> | –105.0 (–318.7, –16.4)<sup>2</sup> | –143.7 (–346.3, –28.1) |

1. Among Jewish children, the difference in estimates of the reduction in odds of carrying +5PCV13 serotypes versus serotype 3, for each PCV13 dose received, is 6.9% (–57.3%, 49.4%).
2. Among Bedouin children, the difference in estimates of the reduction in odds of carrying +5PCV13 serotypes versus serotype 3, for each PCV13 dose received, is 124.4% (21.2%, 322.5%).

Whereas each PCV13 dose conferred a 37.9% (18.2-58.2%) decrease in odds of carrying the six additional serotypes among Jewish children, on average, we identified no association of PCV13 doses received with carriage of these serotypes among Bedouin children (**Table 5**). Distinguishing trends associated with carriage of serotype 3 versus carriage of serotypes 1, 5, 6A, 7F, and 19A, we estimated that Jewish children experienced near-identical reductions of in odds of both outcomes in association with PCV13 doses received. In contrast, Bedouin children experienced 8.1% (–11.5-24.5%) reductions, on average, in odds of carrying serotypes 1, 5, 6A, 7F, and 19A for each PCV13 dose received, but approximately two-fold increases in odds of carrying serotype 3 with each PCV13 dose received.

## DISCUSSION

We report estimates of PCV effectiveness against vaccine-serotype pneumococcal colonization from individually-matched, case-control analyses nested within a large carriage study in southern Israel. We do not identify clear evidence that a single PCV dose protects against vaccine-serotype colonization during the first year of life. While a second dose provides a discernable improvement in protection over the first dose, our point estimates suggest children receiving two doses nonetheless experience only 23-50% protection against vaccine-serotype colonization at ages 5-12 months; our findings do not rule out the possibility that two primary doses are ineffective against vaccine-serotype carriage in the first year of life. In the second year of life and beyond, receipt of a third dose provides a 56-66% improvement in protection against vaccine-serotype colonization, relative to protection sustained after two doses. Our point estimates suggest children receiving all three doses experience roughly 50-77% protection against vaccine-serotype colonization at ages 13-24 months. These findings indicate that, in a setting implementing 2+1 PCV dosing, protection against vaccine-serotype colonization is sustained largely by the third dose. In addition, we identified no evidence of PCV13-derived protection against carriage of serotype 3.

Few studies have directly assessed dose-specific PCV-conferred protection against colonization,^7,18^ and none have addressed post-implementation effectiveness, representing real world protection. In a randomized pre-licensure trial in southern Israel, one and two PCV7 doses were estimated to reduce vaccine-serotype colonization prevalence by –1% (–28-21%) and 27% (2-45%), respectively,^16^ closely resembling our estimates. A meta-analysis identified three studies assessing carriage after one versus zero doses in the primary series.^19^ Consistent with our findings, no studies identified a protective effect of a single PCV dose; across pooled studies, two doses were estimated to confer only 7% (–7-19%) protection against vaccine-serotype colonization.^19^ While our point estimates suggested greater protection after the second dose, our study, too, did not rule out the possibility of 0% two-dose effectiveness at ages 5-12 months.

Elsewhere, 3+0 and a 2+1 series have been estimated to confer 60-69% protection against vaccine-serotype carriage in the second year of life.^20–22^ These findings are in agreement with our point estimates of 50-77% PCV effectiveness against vaccine-serotype carriage after three doses at age 13-24 months.

As Israel uses a 2+1 dosing schedule, it is important to note that our analysis of 3-dose protection in the second year of life generally represents the effect of two primary series doses and a booster dose; low (<1%) three-dose coverage in the first year of life in our study indicates almost all children who had received 3 doses by age 13 months received the third as a booster dose. Our estimates of two-dose protection in the second year of life are not expected to reflect protection under the 1+1 schedule, as children in our study who had received two doses by ages 13-24 months may have received both in the primary series. Although enhanced PCV effectiveness against IPD is associated with receipt of PCV at ages ≥12 months,^9^ it is less clear how age influences vaccine effectiveness against carriage.^7^ It remains critical to determine whether administering a second dose at age 12 months can provide protection comparable to what children experience after receiving three doses under either 2+1 or 3+0 schedules. Recent evidence that indirect protection among adults depends upon the elimination of vaccine-serotype pneumococci among toddlers and school-aged children^23,24^ underscores the need to design schedules maximizing protection among older children.

Whereas we estimate that three PCV7/13 doses confer 74.1% (58.4-84.6%) protection against carriage of PCV7-targeted serotypes in the second year of life, we obtain a non-significantly lower estimate of 49.7% (15.8-83.3%) protection against carriage of all PCV13 serotypes. Ineffectiveness of PCV13 against serotype 3 carriage contributes to this difference. Overall in our study, serotype 3 was the most prevalent of the six serotypes targeted only by PCV13. Receiving more PCV13 doses was associated with reduced odds of carrying serotypes 1, 5, 6A, 7F, and 19A, whereas we did not identify reduced odds of carrying serotype 3 with receipt of more PCV13 doses. Subgroup analyses suggested under-protection against serotype 3 carriage was of particular importance within the Bedouin population; odds of carrying serotype 3 increased in association with PCV13 doses received among Bedouin children. These trends also suggested reduced protection against serotypes 1, 5, 6A, 7F, and 19A among Bedouin children relative to Jewish children, although differences between Bedouin and Jewish children were not otherwise apparent with respect to protection against all PCV13 serotypes or PCV7 serotypes.

The population-level impact of PCVs depends on vaccine-derived protection against colonization. Thus, although the 1+1 schedule may provide non-inferior immunogenicity after the second dose as compared to a 2+1 schedule after the third dose, epidemiologic surveillance reveals that vaccine serotypes, and especially serotypes 3, 14, 19A, and 19F, have not been successfully eliminated from most settings.^25–27^ Because a single primary PCV dose does not prevent carriage acquisition, and offers limited protection against invasive disease,^9^ incidence of vaccine-serotype invasive pneumococcal disease among infants may increase in settings that implement 1+1 PCV schedules. For children receiving two primary doses, we estimate limited and non-significant protection against vaccine-serotype carriage during the first year of life, suggesting that indirect effects observed in countries implementing 2+1 PCV schedules are sustained by protection against vaccine-serotype colonization after the booster dose. Indeed, recent analyses have suggested reductions in vaccine-serotype invasive pneumococcal disease among adults are driven by protection among toddlers and school-aged children, rather than infants.^23^

Several limitations of our study should be considered. Although we do not compare schedules directly, estimates of protection at ages 5-12 months after receipt of one or two doses, and at ages 13-24 months after receipt of three or two doses, may illustrate the relative effects of higher- or lower-dose vaccine series as a substantial proportion of children receive PCV doses off-schedule.^28^ While our study was not randomized, our ability to match children on age, visit timing, prior antibiotic receipt, and ethnicity helps to ensure cases and controls were exposed to a similar risk of carriage acquisition. Drawing both cases and controls from patients in a clinical setting reduces bias that may otherwise result from the association of vaccination with healthcare-seeking behavior.^29^ Statistical power was limited in our study by the fact that there were few unvaccinated children after PCV implementation. Thus, our estimates of the relative effectiveness of two versus one dose, and three versus two doses, convey important information relating to dose-specific protection, as these analyses achieved greater statistical power. Last, it is important to note that our analysis addressed effects of PCVs on prevalence of vaccine-serotype carriage, which may differ from protection against acquisition of vaccine-serotype pneumococci.^30^ Differential prevalence of serotype-specific naturally-acquired immunity among vaccinated and unvaccinated individuals may confound estimates of protection against acquisition.^29^

Randomized assessments of the effect of reduced-dose PCV schedules on pneumococcal colonization are warranted as an increasing number of countries consider schedule changes.^4^ Incorporating cluster randomization, or nasopharyngeal sampling of enrollees’ siblings, would enable such trials to assess how a reduced primary series impacts the indirect effects of PCV use. In addition, transmission-dynamic models parameterized with estimates of vaccine direct effects, such as those we have provided, can yield insight into the potential consequences of reduced-dose schedules in various settings. Such evidence should be sought to understand the impact of reduced-dose schedules on the public health benefits and cost-effectiveness of pediatric PCV programs.

## ACKNOWLEDGMENTS

This study was funded in part by a grant from Pfizer (grant no. 0887X1-4603 to RD). JAL received support from a Robert Austrian Young Investigator award from the International Symposium on Pneumococci and Pneumococcal Diseases.

## SUPPORTING INFORMATION

### CASE-CONTROL SAMPLING AND INFERENCE METHODS

#### Rationale for use of randomly-selected controls

Take *Z*_*i*_=1 to indicate that a child *i* is vaccinated, for *i* in 1, 2, 3, …, *N* children, and take *Z*_*i*_=0 to indicate child *i* is unvaccinated. Define *V* = Σ_*i*_ Pr(*Z*_*i*_ = 1)/*N* as the proportion of children vaccinated. Take *Y*_*i*_=1 to indicate a child carries vaccine-serotype *Streptococcus pneumoniae*, and let *Y*_*i*_=0 indicate the child does not carry vaccine-serotype *Streptococcus pneumoniae* (due to the absence of colonization or carriage of another serotype). Define *P* = Pr(*Y* = 1|*Z* = 0) as the prevalence of colonization among the unvaccinated, and define *θ* as the relative prevalence of vaccine-serotype *Streptococcus pneumoniae* among the vaccinated versus unvaccinated, due only to receipt of the vaccine, such that Pr(*Y* = 1|*Z* = 1) = *θP* and the vaccine effectiveness (VE) is equal to 1 - *θ*.

Under traditional case-control designs, the odds of vaccination among persons who do not experience the outcome against which VE is calculated are assumed to provide a “null” control against which to compare odds of vaccination among persons who experience the outcome. However, in this instance,

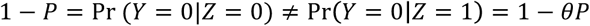

in contrast to the typical assumption. Thus, under the conventional case-control odds ratio (*OR*_*Z*|*Y*_),

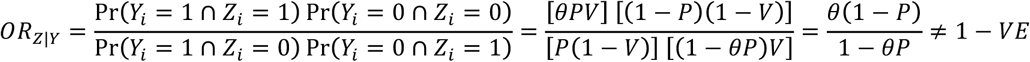

We therefore required an alternative definition of controls under which the relative odds of prior vaccination among cases and controls would reduce to *θ*.

Consider the case where controls are drawn at random from the population, rather than being defined as those individuals for whom *Y=*0:

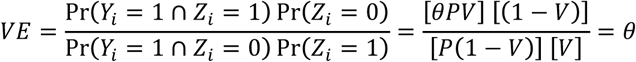

Thus, we selected controls at random from the population rather than defining controls as those with *Y*_*i*_=0.

#### Replacement sampling for case-control inference

Note, however, that this circumstance requires a regime in which selecting cases from the study population does not alter the distribution of *Z* among eligible controls who are not yet sampled. Define *S*_*i*_=1 as an indicator that an individual has been selected as a case, such that Pr(*S*_*i*_ = 1|*Y*_*i*_ = 1) = *C* and Pr(*S*_*i*_ = 1|*Y*_*i*_ = 0) = 0. Under the circumstance where controls are sampled from among those individuals who were not deemed to be cases,

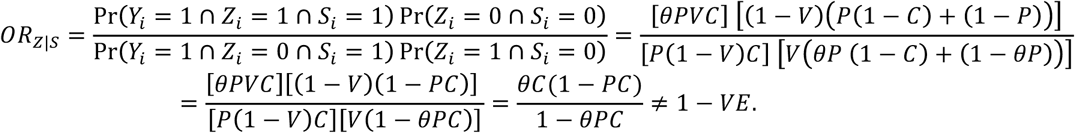

Thus, in order to obtain direction-unbiased VE estimates, we did not exclude those who had been sampled as cases from the eligible control population. Controls, however, were sampled without replacement.

#### Impact on measures of uncertainty

A drawback of this approach is that variance estimates may be artificially reduced if case individuals are re-entered into the analysis as controls. To assess the extent to which this affected our measures of statistical uncertainty, we compared the standard error of the log odds ratio achieved with the re-selected cases to the standard error expected in a scenario where cases were not re-selected as controls, as defined below. Taking *U*_*i*_=1 to indicate an individual was sampled as a control,

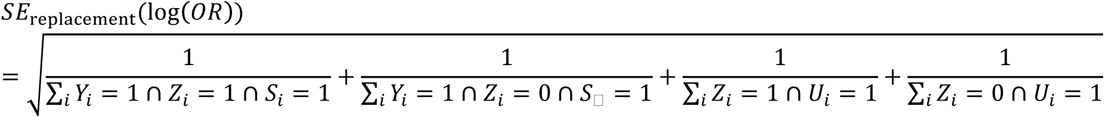

and

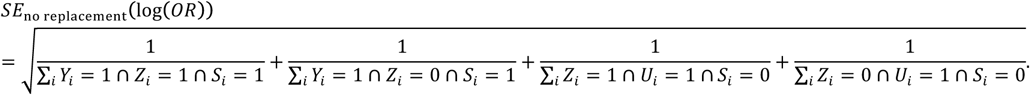

Thus, the ratio 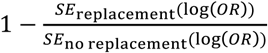 indicates the reduction in width of the confidence intervals expected to result from re-selecting cases as controls.

We report these measures below for analyses of vaccine effectiveness (**Table S1**), together with 95% confidence intervals. Point estimates indicate the standard error was expected to be increased by ≤1%. Thus, retaining case-individuals in the population eligible for selection as controls was not expected to alter measures of uncertainty in any discernable fashion, or to change the conclusions of significance testing.

---

**Table S1:** Degree to which re-selecting cases as controls affects standard error of VE estimates.

| Age (mo.) | Vaccine | Endpoint | Reduction in standard error (95% CI)<br>$1 - \frac{SE_{\text{replacement}}(\log(OR))}{SE_{\text{no replacement}}(\log(OR))}$ |
| --- | --- | --- | --- |
| 5–12 | PCV7/13 only | PCV7 serotypes | 0.2% (0.0%, 1.2%) |
|  | PCV13 only | PCV13 serotypes | 0.6% (0.0%, 2.0%) |
|  |  | +6PCV13 serotypes | 0.3% (0.0%, 1.2%) |
|  | PCV7 only | PCV7 serotypes | 1.0% (0.0%, 3.0%) |
| 13–24 | PCV7/13 only | PCV7 serotypes | 0.1% (0.0%, 0.5%) |
|  | PCV13 only | PCV13 serotypes | 0.1% (0.0%, 0.7%) |
|  | PCV7 only | PCV7 serotypes | 0.1% (0.0%, 0.5%) |
| 13–59 | PCV7/13 only | PCV7 serotypes | 0.2% (0.1%, 0.3%) |
|  | PCV13 only | PCV13 serotypes | 0.1% (0.1%, 0.2%) |
|  | PCV7 only | PCV7 serotypes | 0.1% (0.1%, 0.2%) |

